# Angiotensin-converting enzyme 2 (ACE2) is upregulated in Alzheimer’s disease brain

**DOI:** 10.1101/2020.10.08.331157

**Authors:** Qiyue Ding, Nataliia V. Shults, Brent T. Harris, Yuichiro J. Suzuki

## Abstract

Alzheimer’s disease is a chronic neurodegenerative disorder and represents the main cause of dementia. Currently, the world is suffering from the pandemic of coronavirus disease 2019 (COVID-19) caused by severe acute respiratory syndrome coronavirus 2 (SARS-CoV-2) that uses angiotensin-converting enzyme 2 (ACE2) as a receptor to enter the host cells. In COVID-19, neurological manifestations have been reported to occur. The present study demonstrates that the protein expression level of ACE2 is upregulated in the brain of Alzheimer’s disease patients. The increased ACE2 expression is not age-dependent, suggesting the direct relationship between Alzheimer’s disease and the ACE2 expression. Oxidative stress has been implicated in the pathogenesis of Alzheimer’s disease, and Alzheimer’s disease brains examined in this study also exhibited higher carbonylated proteins as well as increased thiol oxidation state of peroxiredoxin 6 (Prx6). The positive correlation was found between the increased ACE2 protein expression and oxidative stress in Alzheimer’s disease brain. Thus, the present study reveals the relationships between Alzheimer’s disease and ACE2, the receptor for SARS-CoV-2. These results warrant monitoring Alzheimer’s disease patients with COVID-19 carefully for the possible higher viral load in the brain and long-term adverse neurological consequences.

## Introduction

Alzheimer’s disease is a chronic neurodegenerative disorder and the main cause of dementia around the world, affecting a large number of aging populations with an estimated prevalence of 10–30% in the population >65 years of age [Masters et al., 2015; Scheltens et al., 2016]. In the United States, an estimated 5 million people have Alzheimer’s disease, which is the fifth leading cause of death among older adults. However, little is known about the cause of Alzheimer’s disease and no curative treatments are available [Ballard et al., 2011; Querfurth & LaFerla, 2010]. Oxidative stress and the production of reactive oxygen species (ROS) [Freeman & Crapo, 1982; Halliwell & Gutteridge, 2007] have been implicated in Alzheimer’s disease [Tabner et al., 2005; Butterfield et al., 2006; Zhao & Zhao, 2013; Chen & Zhong, 2014]. However, clinical trials on antioxidants have not shown promise as effective treatments [Polidori & Nelles, 2014; Persson et al., 2014]. Thus, further understanding of the role of oxidative stress in Alzheimer’s disease is needed.

Coronaviruses are positive sense single-stranded RNA viruses that often cause the common cold [Su, 2016; Satija, 2007]. Some coronaviruses, however, can be lethal; and currently, the world is suffering from the pandemic of coronavirus disease 2019 (COVID-19) caused by severe acute respiratory syndrome coronavirus 2 (SARS-CoV-2) [Wu, 2020; Huang, 2020]. So far, 30 million people have been infected with SARS-CoV-2 worldwide, causing serious health, economical, and sociological problems. SARS-CoV-2 uses angiotensin-converting enzyme 2 (ACE2) as a receptor to enter the host cells [Yan, 2020; Tai, 2020]. It has been noted that the elderlies, especially those with cardiovascular diseases, is highly susceptible to developing severe COVID-19 conditions [Huang, 2020; Li, 2020; Yang, 2020].

While the current focus is to treat lung and cardiovascular aspects of COVID-19 to reduce the mortality, it has been reported that that neurological manifestations occurred in 36% of COVID-19 patients [Mao, 2020], indicating that this disease may exert to long-term neurological consequences. Thus, understanding the relationships between COVID-19 and the brain is important. The present study monitored the expression level of ACE2, the host cell receptor for SARS-CoV-2, and found that ACE2 protein is upregulated in the brain of Alzheimer’s disease patients.

## Materials and Methods

### Patient samples

The hippocampus is the major structure in the brain affected by Alzheimer’s disease and oxidative stress [Wang & Michaelis, 2010]. Frozen brain (hippocampus) tissues from de-identified deceased patients who had been diagnosed with Alzheimer’s disease and those from control individuals without Alzheimer’s or other neurological diseases were obtained from the Georgetown University Medical Center Brain Bank. Formalin-fixed and paraffin-embedded hippocampus tissues from the same patients were also obtained. Patients usually died at home and full autopsies were not performed on most of these cases. Bodies of deceased patients were maintained in refrigeration until tissue collections. The mean post mortem interval value for Alzheimer’s disease patients was 13.8 ± 1.9 hours and that of controls was 15.2 ± 2.5 hours. After the brain removal, tissues were flash frozen on dry ice and then stored at −80°C. Control brain tissues were obtained using the same protocols. ABC grading for Alzheimer’s disease was performed histologically as previously described [Hyman et al., 2012] at the time of banking by a neuropathologist and co-author, Dr. Brent Harris.

### Tissue homogenate preparations

Frozen hippocampus brain tissues were homogenized in a 4 volume of the 10% trichloroacetic acid (VWR International, Radnor, PA, USA) solution using a Kontes glass tissue grinder. Homogenized tissues were incubated on ice for 30 min and centrifuged for four minutes at 8,000 rpm at 4°C in an accuSpin Micro R centrifuge (ThermoFisher Scientific, Waltham, MA, USA). The resultant pellets were then washed with acetone three times, then with ethanol once. Pellets were air dried, resuspended in the Lysate Buffer (supplied in the -*SulfoBiotics*- Protein Redox State Monitoring Kit Plus, Dojindo Molecular Technologies Inc., Rockville, MD, USA) supplemented with phenylmethylsulfonyl fluoride (2 mM), leupeptin (10 μg/ml), and aprotinin (10 μg/ml) (MilliporeSigma, Burlington, MA, USA) and sonicated on ice using an ultrasonic processor. Samples were then centrifuged at 13,000 rpm for 10 min, and the supernatants were collected. Protein concentrations of cell lysates were measured using the Pierce bicinchoninic acid (BCA) Assay (Thermo Fisher Scientific, Waltham, MA, USA).

### Western blotting

Equal amounts of protein samples (5 μg) were electrophoresed through a reducing sodium dodecyl sulfate polyacrylamide gel. Proteins were then electro-transferred to the Immobilon-FL Transfer Membrane (MilliporeSigma, Burlington, MA, USA). The membranes were blocked with Odyssey blocking buffer (LI-COR, Lincoln, NE, USA) for one hour at 25°C and incubated overnight with the rabbit ACE2 antibody (Catalog # 4355; Cell Signaling), the goat anti-glyceraldehyde 3-phosphate dehydrogenase (GAPDH) antibody (Catalog # PLA0302; MilliporeSigma), or the rabbit anti-peroxiredoxin 6 (Prx6) antibody (Catalog # P0058; MilliporeSigma) at 4°C. Washed membranes were then incubated with IRDye 680RD or IRDye 800CW (LI-COR) for one hour at 25°C in the dark. Signals were obtained by using the Odyssey Infrared Imaging System (LI-COR), and Western blot band intensities were analyzed by densitometry.

### Measurements of protein carbonylation

Carbonyl groups in protein side chains were derivatized with 2,4-dinitrophenylhydrazine (DNPH) [Levine et al., 1994] to form the 2,4-dinitrophenyl (DNP) hydrazone derivative for detection using the OxyBlot Protein Oxidation Detection Kit (MilliporeSigma), according to the manufacturer’s instructions with rabbit anti-DNP antibody at 1:150 dilution for Western blotting. The Immobilon-FL Transfer Membrane was then incubated with anti-rabbit IRDye 680RD for one hour. Signals were then captured by using the Odyssey Infrared Imaging System, and band intensities were analyzed by densitometry.

### Protein thiol redox state monitoring

Protein thiol redox states were monitored using the *-SulfoBiotics-* Protein Redox State Monitoring Kit Plus (Catalog # SB12; Dojindo Molecular Technologies). Samples were labeled with the Protein-SHifter Plus in accordance with the manufacturer’s instructions. After cell extracts were subjected to SDS-PAGE, gels were exposed to the UV light on a transilluminator to remove Protein-SHifter. Proteins in the gel were then electro-transferred to the Immobilon-FL Transfer Membrane. The membrane was blocked with Odyssey Blocking Buffer for one hour at 25°C and incubated overnight with the rabbit anti-Prx6 antibody (Catalog # P0058; MilliporeSigma) at 4°C. Washed membranes were then incubated with anti-rabbit IRDye 680RD for one hour at 25°C in the dark. Signals were then captured by using the Odyssey Infrared Imaging System, and band intensities were analyzed by densitometry.

### Statistical analysis

Means and standard errors of mean (SEM) were computed. Two groups were compared by a two-tailed Student’s *t* test, and differences between more than two groups were determined by the analysis of variance (ANOVA). *P* < 0.05 was defined to be statistically significant.

## Results

### ACE2 protein expression

We performed Western blotting to determine the protein expression levels of ACE2 in hippocampal brain tissues from Alzheimer’s patients and compared with those of control subjects. The ACE2 protein expression was found to be higher in Alzheimer’s brain compared to controls (Fig. 1A). Quantifications of the ratio of ACE2 to GAPDH values revealed that the mean difference was 4.7-fold and statistically significant (Fig. 1B). The mean ACE2 / GAPDH ratio value for 13 Alzheimer’s disease patients was 1.22 ± 0.2, while those of five control subjects were very consistent with a mean of 0.26 ± 0.04. The expression levels of GAPDH were comparable between the control group (21,554 ± 1,021 densitometry units) vs. the Alzheimer’s group (21,315 ± 1,471 densitometry units).

**Figure 1:**
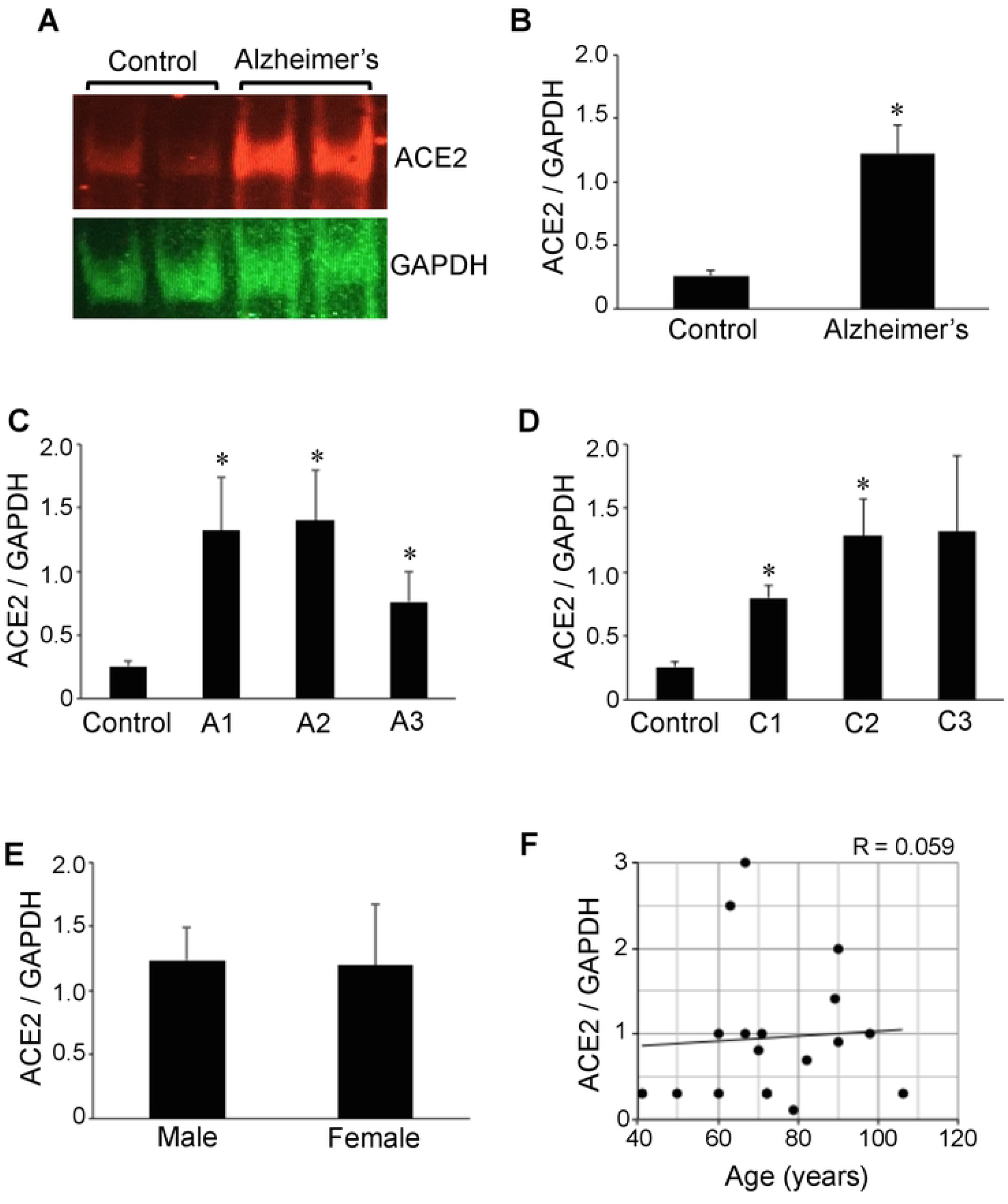
The ACE2 protein expression is upregulated in brains of Alzheimer’s disease patients. Brain (hippocampus) homogenates from control subjects and Alzheimer’s disease patients were subjected to Western blotting using antibodies against ACE2 and GAPDH. (A) Representative results. (B) The bar graph represents means ± SEM of the ratio of ACE2 to G3PDH (N = 13 for Alzheimer’s and 5 for Control). (C) The bar graph represents means ± SEM of the ratio of ACE2 to G3PDH for various Aβ plaque scores (denoted by the letter A). N = 5 for Control, 5 for A1, 5 for A2 and 3 for A3. (D) The bar graph represents means ± SEM of the ratio of ACE2 to G3PDH for various neuritic plaque score (denoted by the letter C). N = 5 for Control, 2 for C1, 7 for C2 and 4 for C3. The symbol * denotes that the value is significantly different from the control value at *P*< 0.05. (E) The bar graph represents means ± SEM of the ratio of ACE2 to G3PDH (N = 8 for male and 5 for female). No significant difference was found between males and females. (F) The scattered graph represents age vs. ACE2 expression (N = 18). Pearson correlation coefficient R = 0.059 (weak positive correlation).

The severity of Alzheimer’s patients examined in this study was mostly in the intermediate to high range with 61.5% of patient samples exhibiting Aβ plaque scores of 2 or 3 and 84.6% exhibiting neuritic plaque scores of 2 or 3. The analysis of ACE2 protein expression in relation to Aβ plaque scores (Fig. 1C) or neuritic plaque scores (Fig. 1D) determined that the ACE2 expression levels are not related to the severity of the disease and the significant upregulation of ACE2 occurs even in Alzheimer’s disease patients with low severity. Further, among the Alzheimer’s disease patients, no gender difference in ACE2 levels was noted (Fig. 1E). While, in the present study, the age of Alzheimer’s disease patients ranged from 60 to 106 (mean = 78.8 ± 4.2 years) and that of control subjects ranged from 41 to 79 (mean = 60.4 ± 7.8), our analysis detected no age-dependence in the ACE2 protein expression (Fig. 1F).

Silver staining images of our formalin-fixed paraffin-embedded tissues from Alzheimer’s patients showed amyloid and neuritic plaques as well as the neurofibrillary tangles [Hedreen et al., 1994; Reusche, 1991]. Further, hematoxylin and eosin staining detected the amyloid plaques [DeTure & Dickson, 2019; Ryan et al., 2015; Serrano-Pozo et al., 2011] and immunohistochemistry using the Tau AT8 antibody demonstrated the expression of phosphorylated Tau at serine 202 and threonine 205 [Goedert et al., 1995] in our hippocampal tissue samples from Alzheimer’s disease patients (data not shown).

### Protein carbonylation

Oxidative stress has been shown to occur in Alzheimer’s disease brain [Tabner et al., 2005; Butterfield et al., 2006; Zhao & Zhao, 2013; Chen & Zhong, 2014]. Consistently, protein carbonylation of various hippocampal proteins as monitored by the DNPH derivatization of carbonylated proteins using the OxyBlot Kit showed higher carbonylated proteins in Alzheimer’s brain compared controls (Fig. 2A). Quantifications of the results revealed that the protein carbonyl content is 7.8-fold higher in Alzheimer’s disease patients compared to controls and the difference was found to be statistically significant (Fig. 2B). The carbonyl content-to-GAPDH ratio values ranged from 0.7 to 2.9 (mean = 1.4 ± 0.15) in Alzheimer’s disease brain while controls had a range of 0.1 to 0.2 (mean = 0.2 ± 0.02). Our analysis of Aβ plaque scores (Fig. 2C) or neuritic plaque scores (Fig. 2D) in relation to hippocampal brain protein carbonyl content may have shown a tendency to be dependent on the severity of Alzheimer’s disease, but no significant differences were noted among patient brain tissues. The protein carbonyl content was found to be consistently higher with statistical significance between Alzheimer’s patients of any degree of severity and controls.

**Figure 2:**
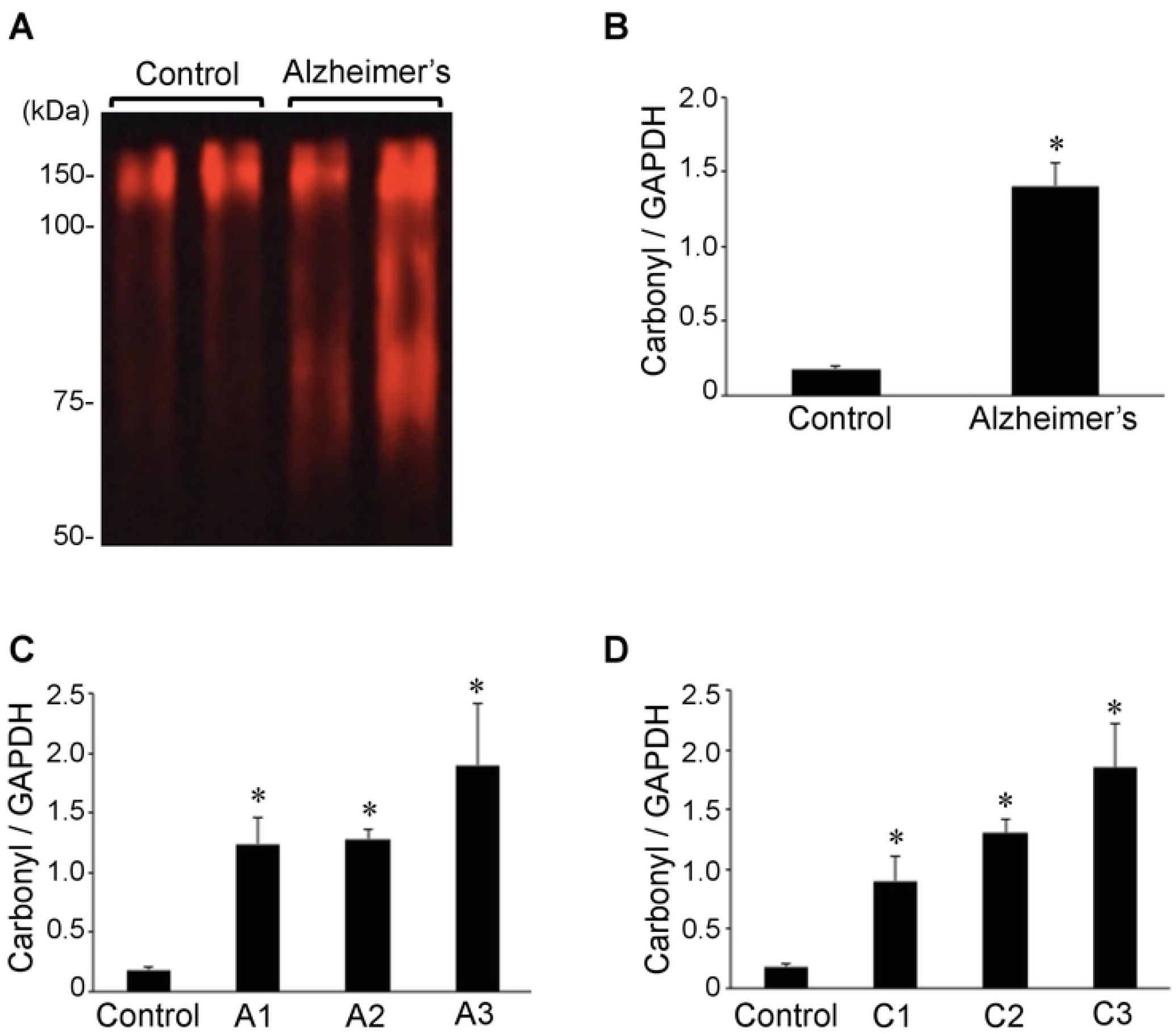
Protein carbonylation is upregulated in brains of Alzheimer’s disease patients. Brain (hippocampus) homogenates from control subjects and Alzheimer’s patients were studied using OxyBlot. (A) Representative results. (B) The bar graph represents means ± SEM of the ratio of carbonylated proteins to G3PDH (N = 13 for Alzheimer’s and 5 for Control). (C) The bar graph represents means ± SEM of the ratio of carbonylated proteins to G3PDH for various Aβ plaque scores (denoted by the letter A). N = 5 for Control, 5 for A1, 5 for A2 and 3 for A3. (D) The bar graph represents means ± SEM of the ratio of carbonylated proteins to G3PDH for various neuritic plaque score (denoted by the letter C). N = 5 for Control, 2 for C1, 7 for C2 and 4 for C3. The symbol * denotes that the value is significantly different from the control value at *P*< 0.05.

### Protein Redox State Monitoring Kit Plus analysis of Prx6

The *-SulfoBiotics-* Protein Redox State Monitoring Kit Plus allows to assess thiol redox status of a given protein by using 15-kDa Protein-SHifters that are added to the reduced cysteine sulfhydryl groups followed by immunological detection of the proteins of interest [Suzuki et al., 2019]. In case of an antioxidant, human Prx6, that contains two cysteines, when both cysteine residues are oxidized, no Protein-SHifter bind to the protein, thus a 25-kDa band that is the molecular weight of Prx6 is observed. If one cysteine is reduced, then one 15-kDa Protein-SHifter binds to this cysteine, forming a 40-kDa complex, shifting the Prx6 band. If both cysteine residues are reduced, two Protein-SHifters bind to the protein allowing a total of 30-kDa shift of the 25-kDa Prx6, forming a 55-kDa species. We have previously found that, in untreated cultured human smooth muscle cells, Prx6 molecules are predominant in the reduced state, exhibiting a 55-kDa band in the *-SulfoBiotics-* Protein Redox State Monitoring analysis [Suzuki et al., 2019]. The treatment of cells with hydrogen peroxide resulted in the conversion of the 55- kDa band into mostly 40-kDa, rather than 25-kDa, suggesting that only one of the two cysteines in human Prx6 protein is susceptible for this oxidation [Suzuki et al., 2019]. Series of experiments determined that the shift of 55-kDa to 40-kDa species is due to the oxidation of the catalytic cysteine 47 [Suzuki et al., 2019].

Using this system, we monitored protein thiol status in Alzheimer’s hippocampal brains and controls. Similarly to the situation that can be produced by the treatment of cultured cells with hydrogen peroxide [Suzuki et al., 2019], we found that a large portion of Prx6 protein molecules in the Alzheimer’s brain exist as the 40-kDa species that corresponds to the addition of one Protein-SHifter (Fig. 3A). Densitometry analysis determined that the ratio of 40-kDa to 55-kDa Prx6 species is 2.5-fold higher in Alzheimer’s brain compared to control (Fig. 3B), suggesting that catalytic cysteine is preferentially oxidized in the Alzheimer’s brain. By contrast, no significant differences were noted for either the 25-kDa to 55-kDa or 25-kDa to 40-kDa ratio (data not shown), indicating that non-catalytic cysteine is not consistently oxidized in the Alzheimer’s disease brain. The range of the ratio of 40-kDa to 55-kDa Prx6 species in Alzheimer’s disease patients was from 0.3 to 1.2 (mean = 0.9 ± 0.08), while that of controls was from 0.2 to 0.5 (mean = 0.36 ± 0.05). Prx6 thiol oxidation as measured by the ratio of 40-kDa to 55-kDa species was found to correlate with protein carbonylation (Fig. 3C).

**Figure 3:**
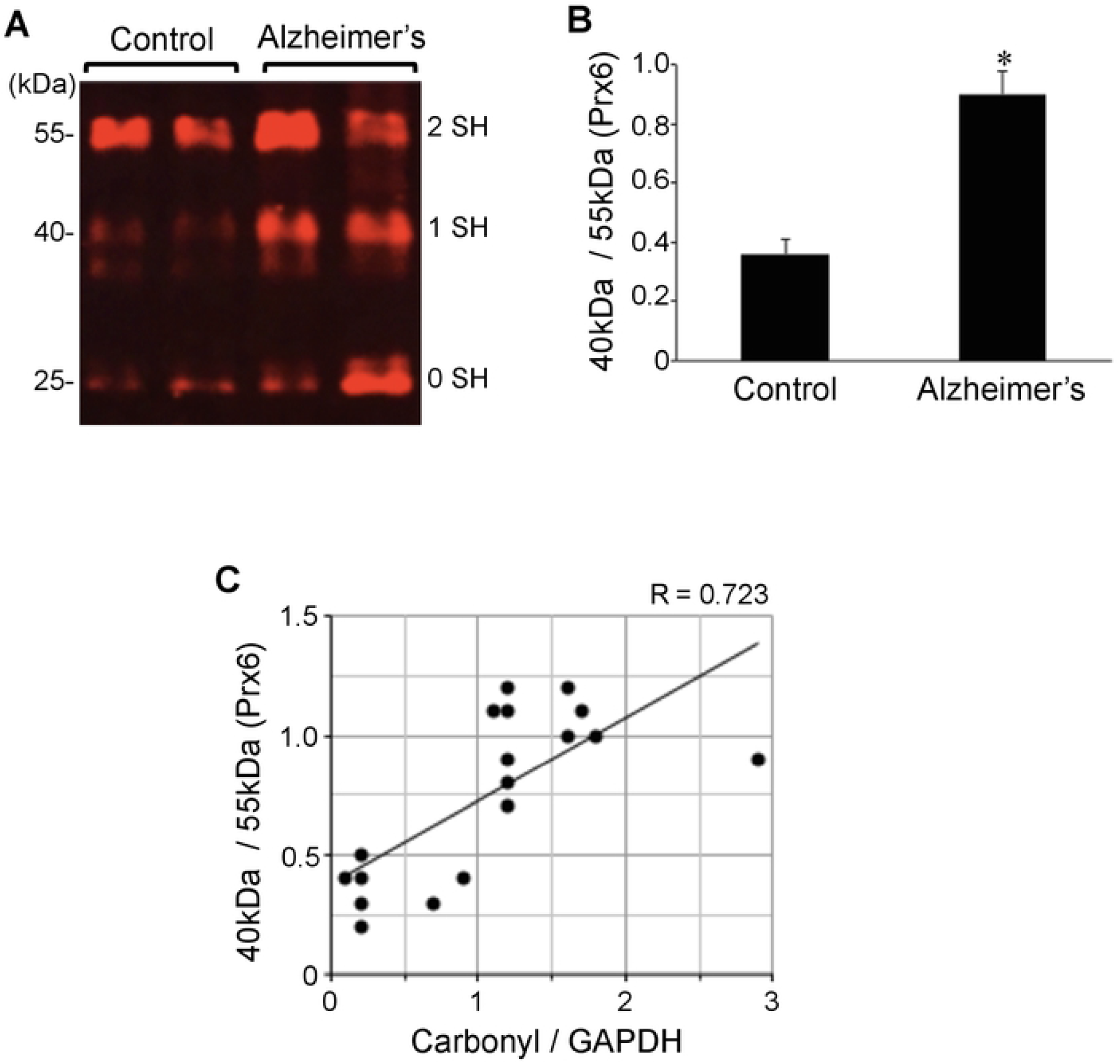
Cysteine oxidation of Prx6 is upregulated in brains of Alzheimer’s disease patients. Brain (hippocampus) homogenates from control subjects and Alzheimer’s patients were investigated for cysteine oxidation of Prx6 using - *SulfoBiotics* - Protein Redox State Monitoring Kit Plus. A free protein thiol group was labeled with the Protein-SHifter Plus that contains maleimide with high affinity toward reduced sulfhydryl groups. Each Protein-SHifter causes a 15-kDa shift. After electrophoresis, the Protein-SHifter Plus moiety was eliminated by the exposure of the gel to ultraviolet light that increases the efficiency of Western blotting and allowing for detection of specific proteins in biological samples. In case of human Prx6, the protein with fully oxidized cysteine residues migrates at 25 kDa. The protein with one cysteine reduced binds to a Protein-SHifter and migrates at 40 kDa, and the protein with two reduced cysteine residues migrates at 55 kDa. Our previous study determined that the 40-kDa band depicts Prx6, in which the catalytic cysteine (Cys47) is oxidized [Suzuki et al., 2019]. (A) Representative Western blotting results using the Prx6 antibody showing 25-kDa (0 reduced sulfhydryl), 40-kDa (1 reduced sulfhydryl), and 55-kDa (2 reduced sulfhydryl) bands. (B) The bar graph represents means ± SEM of the ratio of 40-kDa Prx6 band to 55-kDa Prx6 band (N = 13 for Alzheimer’s and 5 for Control). (C) The scattered graph represents protein carbonylation vs. Prx6 thiol oxidation (N = 18). Pearson correlation coefficient R = 0.723 (positive correlation).

The analysis of the Aβ plaque or neuritic plaque scores in relation to the ratio of 40-kDa to 55-kDa Prx6 species showed no statistically significant differences noted among patients brain tissues, while the Prx6 thiol oxidation was consistently higher with statistical significance between Alzheimer’s patients of any degree of severity and controls (data not shown).

### Prx6 protein expression

We also found that the expression level of the Prx6 protein is higher in the brain of Alzheimer’s disease patients compared to controls (Fig. 4A), which may reflect the activation of an antioxidant defense mechanism in response to oxidative stress. The use of GAPDH protein as a loading control and the densitometry analysis showed that the protein expression levels of Prx6 between Alzheimer’s and control brains are significantly (6.5-fold) different (Fig. 4B). The range of the Prx6 to GAPDH ratio determined in Alzheimer’s patients’ brains was from 0.9 to 2.5 (mean = 1.4 ± 0.1), while that of control brain was from 0.01 to 0.5 (mean = 0.2 ± 0.08). The Prx6 protein expression was found to correlate strongly with protein carbonylation (Fig. 4C) as well as the Prx6 thiol oxidation as measured by the ratio of 40-kDa to 55-kDa species (Fig. 4D).

**Figure 4:**
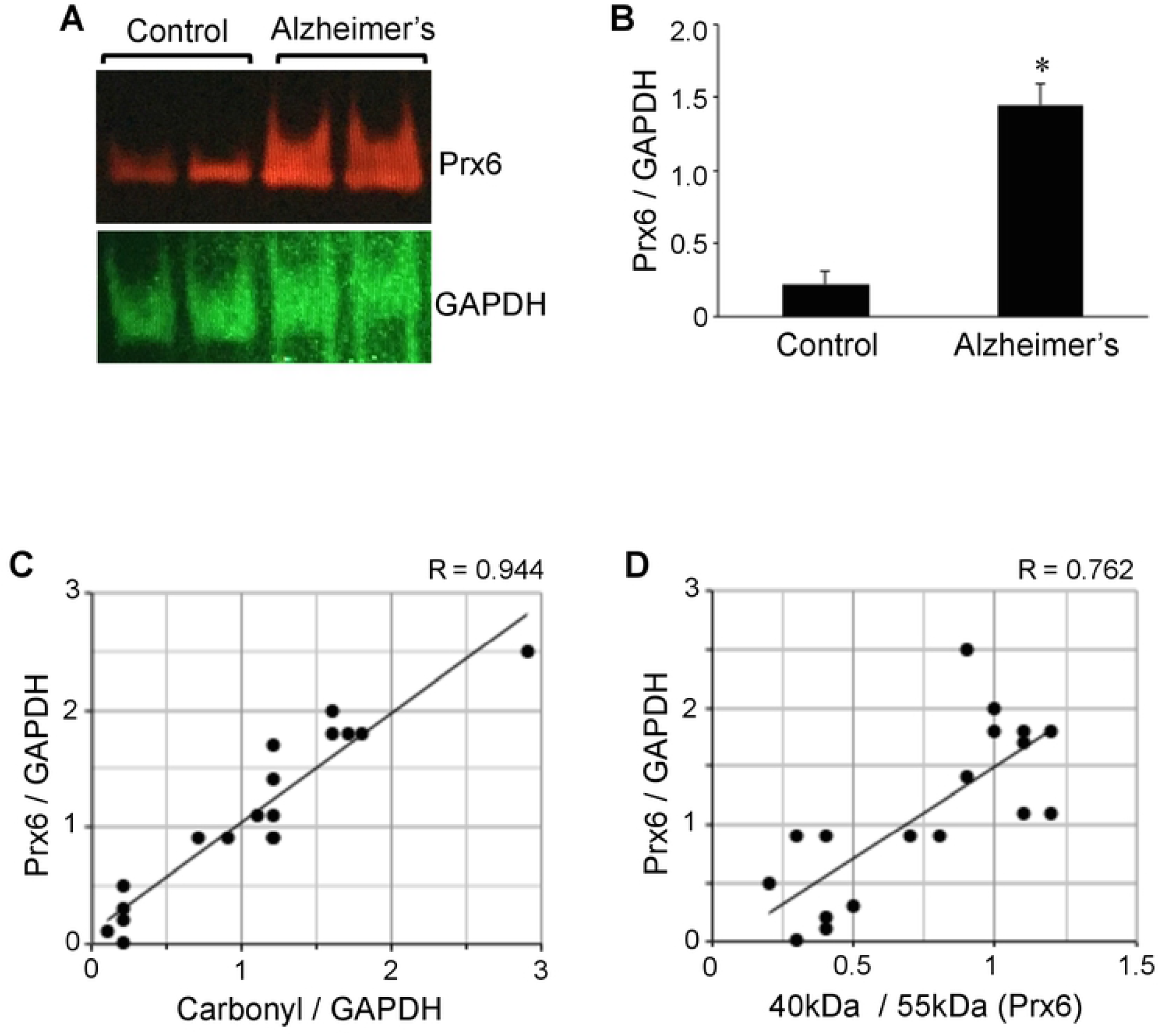
The Prx6 protein expression is upregulated in brains of Alzheimer’s disease patients. Brain (hippocampus) homogenates from control subjects and Alzheimer’s disease patients were subjected to Western blotting using antibodies against Prx6 and GAPDH. (A) Representative results. (B) The bar graph represents means ± SEM of the ratio of Prx6 to G3PDH. The symbol * denotes that the value is significantly different from the control value at *P*< 0.05. (C) The scattered graph represents protein carbonylation vs. Prx6 expression (N = 18). Pearson correlation coefficient R = 0.944 (strong positive correlation). (D) The scattered graph represents Prx6 thiol oxidation vs. Prx6 expression (N = 18). Pearson correlation coefficient R = 0.762 (strong positive correlation).

The analysis of Aβ plaque and neuritic plaque scores in relation to hippocampal brain Prx6 levels may have shown a tendency to be dependent on the severity of Alzheimer’s disease, but no statistically significant differences were noted among patients brain tissues. The Prx6 content was found to be consistently higher with statistical significance between Alzheimer’s patients of any degree of severity and controls (data not shown).

### Correlation analysis to examine the relationships between ACE2 and oxidative stress

Fig. 6 shows the results of correlation analyses to examine the relationships between ACE2 and various oxidative stress parameters using all of the 18 samples (13 Alzheimer’s disease patients and 5 controls). The Pearson correlation coefficient R-value between the ACE2 expression and the protein carbonyl content was found to be 0.517 (Fig. 5A), and the R-value between the ACE2 expression and the Prx6 thiol oxidation state was 0.567 (Fig. 5B). The R-value between the ACE2 expression and the Prx6 expression was found to be slightly higher with 0.687 (Fig. 5C). Thus, the ACE2 protein expression has moderate positive correlation with oxidative stress parameters and the upregulation of an antioxidant.

**Figure 5:**
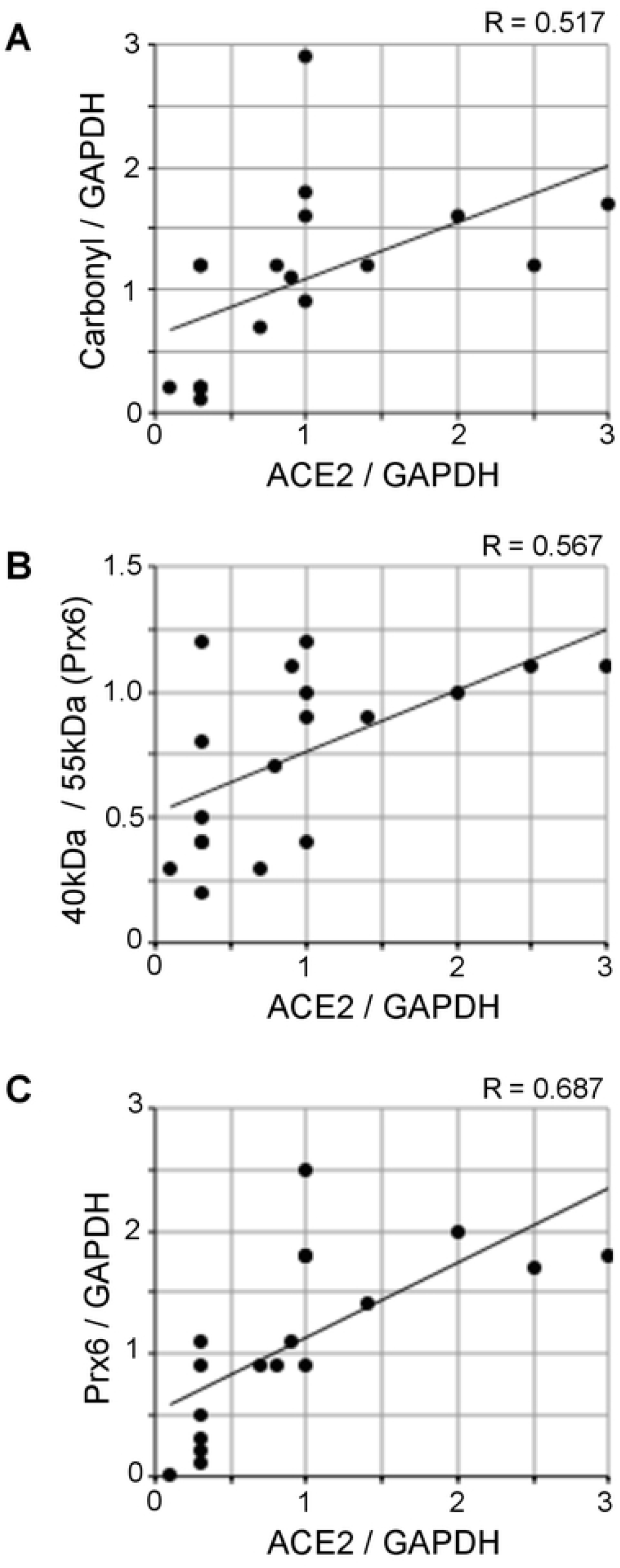
Correlation analysis to examine the relationship between ACE2 protein expression and oxidative stress. The scattered graphs represent (A) ACE2 expression vs. protein carbonylation, (B) ACE2 expression vs. Prx6 thiol oxidation, and (C) ACE2 expression vs. Prx6 expression using Western blotting data (N = 18). Pearson correlation coefficients indicate moderate positive correlations.

## Discussion

The major physiological function of ACE2 is to lower blood pressure by catalyzing the hydrolysis of angiotensin II, which acts as a vasoconstrictor, into angiotensin (1-7) that functions as a vasodilator [Gheblawi et al., 2020]. ACE2 also serves as the receptor for SARS-CoV-2 that is causing the current pandemic of COVID-19. Alveolar epithelial cells and cells of the conducting airways are the primary targets of SARS-CoV-2, resulting in severe pneumonia and acute respiratory distress syndrome (ARDS) [Xu, 2020]. In the brain, immunohistochemistry detected the ACE2 protein expression in endothelial and arterial smooth muscle cells [Hamming et al., 2004]. In addition, cultured brain glial cells [Gallagher et al. 2006] and mouse brain neurons [Doobay et al. 2007] have been shown to express ACE2 [Xia & Lazartigues, 2008].

The major finding of the present study is that the brain ACE2 expression is higher in Alzheimer’s disease patients compared to controls. We did not detect correlation between the severity of Alzheimer’s disease and the ACE2 expression, and the ACE2 was found to be upregulated even in the mild Alzheimer’s disease cases. These results indicate that the SARS-CoV-2 infection of Alzheimer’s disease patients with any disease severity may results on the higher viral entry into the cells in the brain and Alzheimer’s disease patients may be highly affected by COVID-19. No gender difference was noted in the ACE protein expression in Alzheimer’s brain, and the ACE2 protein expression was not dependent on age. Thus, it appears that there is a direct relationship between Alzheimer’s disease and the ACE2 expression. Understanding the mechanism of this relationship may shed a light in the pathogenesis of Alzheimer’s disease as well as possible management strategies for individuals infected with SARS-CoV-2 since ACE2 is the host cell receptor for this virus.

Oxidative stress has been implicated in Alzheimer’s disease and there have been many studies monitoring various oxidative stress parameters [Tabner et al., 2005; Butterfield et al., 2006; Zhao & Zhao, 2013; Chen & Zhong, 2014]. Thus, redox processes may play a viral role in regulating the ACE2 gene expression. Consistent with the previous study [Aksenov et al., 2001], DNPH-reactive protein carbonyls were found to be increased in our Alzheimer’s disease brain samples.

Information about protein thiol oxidation in human Alzheimer’s brain has been limited. Thiol redox status of proteins plays pivotal roles in biology [Boronat & Domènech, 2017; Chung et al., 2013; Sen, 2000; Suzuki et al., 1997]. Since proteins are the functional molecules in the biological system, their redox status directly reflects pathophysiology. While there are a number of ways to monitor the oxidation of sulfhydryl groups in the biological samples, the most of the techniques only allow for the global assessment of the oxidation in cell or tissue samples [Rudyk & Eaton, 2014] and the monitoring of the redox status of specific proteins cannot be readily performed. Some techniques can identify individual proteins that are oxidized, but these approaches are difficult, time consuming, and expensive [Leichert et al., 2008; Couvertier et al., 2014]. The - *SulfoBiotics* - Protein Redox State Monitoring Kit Plus is remarkable in that it can readily allow for the assessment of the redox states of specific proteins of interest in the biological samples in a convenient and cost-effective fashion [Suzuki et al., 2019]. To understand the thiol biology, we previously performed a redox state monitoring analysis of Prx6 using the *-SulfoBiotics-*Protein Redox State Monitoring Kit Plus system [Suzuki et al., 2019]. In this system, a 15-kDa Protein-SHifter is added to every reduced cysteine residue, defining protein thiol redox states while examining the mobility shift caused by the Protein-SHifters using gel electrophoresis. Thiol status specifically on Prx6 was subsequently determined by Western blotting. In human, the Prx6 protein molecule contains one catalytic cysteine and one additional non-catalytic cysteine. We found that the treatment of cultured cells with hydrogen peroxide caused the oxidation of the catalytic cysteine with a minimal influence on the non-catalytic cysteine [Suzuki et al., 2019]. In Alzheimer’s disease patient brain samples, the present study also showed that the catalytic cysteine is preferentially oxidized without significant influence on the non-catalytic cysteine of Prx6. The protein thiol oxidation through the measurement of the ratio of 40-kDa to 55-kDa species of Prx6 correlated with the degree of protein carbonylation.

Peroxiredoxins are a class of peroxidase antioxidant enzymes [Rhee, 2016; Rhee & Kil, 2017]. Six members of peroxiredoxins have been identified [Kim et al., 1988; Seo et al., 2000; Rhee, 2016; Rhee & Kil, 2017]. Peroxiredoxins 1 – 5 contain two catalytic cysteines that perform the two-electron reduction of hydrogen peroxide to water [Rhee, 2016; Rhee & Kil, 2017]. Prx6 is a unique peroxiredoxin that possesses only one catalytic cysteine and presumably utilizes glutathione to receive the second electron for the two-electron reduction [Fisher, 2011; Fisher, 2017]. The present study also found that the Prx6 protein expression is higher in brains of Alzheimer’s disease patients, supporting the concept that these brains have been adapted to contain increased antioxidant defenses. Similarly to our results on Prx6, it has been reported that the protein expression levels of peroxiredoxin 1 and 2 in the brain are also increased in Alzheimer’s disease [Kim et al., 2001]. In our study, the increased Prx6 protein expression was strongly correlated with the degree of oxidative stress as measured by protein carbonylation and Prx6 thiol oxidation.

The correlation analysis of the brain ACE2 protein expression and various oxidative stress parameters including protein carbonylation, cysteine oxidation of Prx6, and Prx6 protein expression showed positive relationships. The relationship between ACE2 and Prx6 was found to be the highest. However, the correlations between these redox parameters and the ACE2 expression were found to be modest, suggesting other factors may also participate in regulating ACE2 gene expression in Alzheimer’s disease. Whether the mechanism of the upregulation of ACE2 expression involves oxidative stress and whether increased ACE2 promotes oxidative stress need further investigations.

In summary, the examination of brains from Alzheimer’s disease patients indicated that ACE2 protein is upregulated in association with oxidative stress. Further work is needed to determine whether redox processes participate in the mechanism of ACE2 gene expression or whether ACE2 regulates oxidative stress in the brain. We analyzed 13 Alzheimer’s disease patients and 5 controls. While the number of control subjects is relatively small, observed values for these individuals were very consistent in all the parameters measured in the present study. Also, in our study, the mean age was higher for Alzheimer’s disease patients than for controls, however, the ACE2 protein expression was found not to be dependent on age with no correlation detected (with a very low Pearson correlation coefficient of 0.059). Thus, it is not likely that these limitations hamper our conclusion that Alzheimer’s disease per se is associated with increased ACE2 protein expression. Our conclusion is further supported by a communication by Lim et al., [2020] published as a Letter to the Editor that their genome-wide association study detected upregulated *Ace2* mRNA in the hippocampus of Alzheimer’s disease patients. In light of the importance of ACE2 in the current pandemic, further investigations on the issue of ACE2 in Alzheimer’s disease brain is warranted, considering the possibility that the high level of ACE2 in Alzheimer’s disease patients may affect their responses to COVD-19.

